# Characterization of Mice Bearing Humanized Androgen Receptor Genes (h/mAr) Varying in Polymorphism Length

**DOI:** 10.1101/2020.08.18.248153

**Authors:** Zsuzsa Lindenmaier, Yohan Yee, Adrienne Kinman, Darren Fernandes, Jacob Ellegood, Christie L. Burton, Diane M. Robins, Armin Raznahan, Paul Arnold, Jason P. Lerch

**Affiliations:** Mouse Imaging Centre, Hospital for Sick Children, Toronto, Ontario, Canada; Department of Medical Biophysics, University of Toronto, Toronto, Ontario, Canada; Psychiatry, Neurosciences and Mental Health, The Hospital for Sick Children, Toronto, Ontario, Canada; Department of Human Genetics, University of Michigan Medical School, Ann Arbor, Michigan, USA; Developmental Neurogenomics Unit, National Institute of Mental Health, Bethesda, Maryland, USA; Department of Psychiatry, University of Calgary, Calgary, Alberta, Canada; Wellcome Centre for Integrative Neuroimaging (WIN), FMRIB, Nuffield Department of Clinical Neuroscience, University of Oxford, Oxford, UK

## Abstract

The androgen receptor (AR) is known for masculinization of behavior and brain. To better understand the role that AR plays, mice bearing humanized *Ar* genes with varying lengths of a polymorphic N-terminal glutamine (Q) tract were created (Albertelli et al 2006). The length of the Q tract is inversely proporitional to AR activity. Biological studies of the Q tract length may also provide a window into potential AR contributions to sex-biases in disease risk.

Here we take a multi-pronged approach to characterizing AR signaling effects on brain and behavior in mice using the humanized *Ar* Q tract model. We first map effects of Q tract length on regional brain anatomy, and consider if these are modified by gonadal sex. We then test the notion that spatial patterns of anatomical variation related to Q tract length could be organized by intrinsic spatiotemporal patterning of AR gene expression in the mouse brain. Finally, we test influences of Q tract length on four behavioral tests.

Altering Q tract length led to neuroanatomical differences in a non-linear dosage-dependent fashion. Gene expression analyses indicated that adult neuroanatomical changes due to Q tract length are only associated with neurodevelopment (as opposed to adulthood). No significant effect of Q tract length was found on the behavior of the three mouse models. These results indicate that AR activity differentially mediates neuroanatomy and behavior, that AR activity alone does not mediate sex differences, and that neurodevelopmental processes are associated with spatial patterns of volume changes due to Q tract length in adulthood. They also indicate that androgen sensitivity in adulthood does not directly lead to autism-related behaviors or neuroanatomy, although neurodevelopmental processes may play a role earlier. Further study into sex differences, development, other behaviors, and other sex-specific mechanisms are needed to better understand AR sensitivity, neurodevelopmental disorders, and the sex difference in their prevalence.

## 1 Introduction

In mammals, the androgen receptor (AR), a ligand-activated transcription factor, is responsible for male primary and secondary sexual differentiation. AR also influences numerous physiological processes not directly linked to reproduction,^1^ like mediating the effect of testosterone on white matter,^2^ modulating intra-abdominal adiposity and blood pressure,^3^ and homeostasis and tumorigenesis of the prostate.^1^

ARs masculinize behavior and brain, including regions like the locus coeruleus and the ventromedial hypothalamus, and behaviors like stress response and cognitive processing (see Zuloaga et al 2008 for review).^4^ In turn, genetic variation in *Ar* may contribute to disease risk and progression.^5^ One of the most studied polymorphisms in the human AR is a variation in the length of a CAG trinucleotide repeat, encoding a polyglutamine (Q) tract within the N-terminal domain. In the general population, Q tract length ranges from 9 to 37 residues, with most having lengths in the 15-30 range.^5^ Q tract length is inversely proportional to AR activity and has been shown to influence prostate cancer progression and oncogenesis,^5^ white and gray matter volume,^2, 6^ body fat mass,^7^ high-density lipoprotein cholesterol levels,^8^ and others.

To better elucidate the role of the Q tract lengths in a controlled manner, Albertelli et al 2006^1^ generated three “humanized” AR mouse lines: a short 12Q, a median 21Q, and a long 48Q. Given that the length of the Q tract is inversely proportional to AR activity, the 12Q (short) mice had increased AR activity, the 48Q (long) had decreased AR activity, and the 21Q mice were in between. Given that some differences exist in the *Ar* gene of the mouse genome,^9^ humanizing the gene allows for minimization of variation and enhanced relevance to human studies.^1^ The generated mice grossly appeared normal in growth, behavior, fertility and reproductive tract morphology, while showing subtle differences in body fat amount and seminal vesicle weight.^1^

Biological studies of the *Ar* CAG repeat polymorphism may also provide a window into potential *Ar* contributions to sex-biases in disease risk. For example, autism spectrum disorder (ASD) is highly male-biased in its prevalence,^10^ and several studies have explored the potential role of androgen signaling in shaping the increased risk in males.^11–18^ For example, as detailed in Henningsson et al 2009,^18^ direct and indirect elevated fetal testosterone levels are associated with autism and related traits,^12–15^ and girls with congential adrenal hyperplasia (associated with high prenatal androgens) display autism-like traits to a higher extent than their unaffected siblings.^16^ Studies in adults show preliminary support for the association of autistic traits with enhanced serum levels of testostrone.^11, 17^ One study of a sample of 267 subjects with ASD even found a higher prevalence of short Q tracts in cases versus control.^18^ Although much data suggests a role of androgens in the disorder, the exact underpinnings remains to be investigated.

Here we take a multi-pronged approach to characterising AR signaling effects on brain and behavior in mice using the humanized *Ar* Q tract model. We first map effects of Q tract length on regional brain anatomy, and consider if these are modified by gonadal sex. We then test the notion that spatial patterns of anatomical variation related to Q tract length could be organized by intrinsic spatiotemporal patterning of *Ar* gene expression in the mouse brain. Finally, we test influences of Q tract length on four behavioral tests.

## 2 Methods

### 2.1 Subjects

Mice bearing humanized *Ar* genes (*h/mAr*) with varying lengths of a polymorphic N terminal glutamine (Q) tract were created (see Albertelli et al 2006^1^). Two cohorts of adult mice, with three varying lengths (and wildtype (Wt) controls), were used. One cohort was naive and sacrificed around p65 for *ex vivo* magnetic resonance imaging (MRI) and involved hemizygous males and homozygous females (the *Ar* gene is on the X chromosome). The other cohort underwent four different behavioral tests, as well as MR imaging at p65, but focused only on hemizygous male mice. All mice were on the C57BL/6J (JAX #000664) background.

Active colonies for each strain were maintained at The Centre for Phenogenomics (TCP). All animals were in temperature-controlled housing, with a 12-hour light-dark cycle, and water and food were available *ad libitum*. Breeders were set up to yield control litter-mates. Mice were group housed, and an attempt was made to achieve a sample size of at least 10 per each group (see Table 1). In cohort 2, data was collected and analyzed for all mice for all behavioral tests. All procedures were approved by the Institutional Animal Care and Use Committee at TCP.

**Table 1:** Tables depicting the sample size in each group, broken down by cohort. A minimum of n=10 was aimed for per genotype, per sex. Table (a) shows the samples for the naive mice that only underwent *ex vivo* imaging around p65. Table (b) shows the cohort that was subjected to behavioral tests along with imaging around p65. Wt= wildtype

| (a) Cohort 1: Naive |  |  | (b) Cohort 2: Behavior |  |
| --- | --- | --- | --- | --- |
| Strain-Genotype | Males | Females | Strain-Genotype | Males |
| Wt | 25 | 10 | Wt | 22 |
| 12Q Hem/Hom | 24 | 21 | 12Q Hem | 14 |
| 21Q Hem/Hom | 17 | 15 | 21Q Hem | 17 |
| 48Q Hem/Hom | 21 | 18 | 48Q Hem | 11 |

### 2.2 Neuroanatomical Phenotype

One *ex vivo* MRI scan was performed at least a month after the mice were sacrificed around postnatal day 65.^19^ The procedure closely follows the procedure described in Lerch et al 2011^20^ and Nieman et al 2018.^21^ Briefly, to prepare the brains for scanning and enhance contrast, a fixation and perfusion procedure was done as previously described in Cahill et al 2012.^22^ A multi-channel 7.0T magnet (Agilent Technologies, Santa Clara, CA) was used, with the ability to scan up to 16 brains simultaneously. A T2-weighted 3D fast spin echo sequence was used to yield an isotropic resolution of 40 *u*m (cylindrical acquisition, TR=350ms, TE=12ms, ~14 hour scan time).^23^

Imaging data was analyzed using image registration and deformation based morphometry approaches as described in Lerch et al (2011).^20^ Briefly, images were linearly (6 parameter, then 12 parameter) and nonlinearly registered together. At completion of this registration, all scans were deformed into alignment with each other in an unbiased fashion. These registrations were performed with a combination of `mni_autoreg` tools^24^ and ANTS (advanced normalization tools).^25, 26^ The changes within regions and across the brain were examined using deformation based morphometry (DBM) and MAGeT (a multi-atlas registration-based segmentation tool)^27^ to calculate the volume of segmented brain structures as well as DBM to calculate any voxel-wise differences. An atlas that segments 182 structures in the adult mouse brain was used.^28–32^ Segmentations of all brains passed quality control by visual inspection.

Imaging data was then analyzed using the R statistical language (R Core Team, 2017).^33^ The data from all mice was first assessed using a linear model to determine the effect of sex, strain, and their interaction, while covarying for cohort (see Equation 1). Subsequently, the wildtype (Wt) mice were removed from the analysis and two models were fit where strain was treated as a factor (dummy coded, referred to as “linear”)(Equation 1) and as an ordered factor (polynomial coded, referred to as “quadratic”)(Equation 2), to assess the effect of Q tract length on neuroanatomy. Additionally, relative brain region volumes were calculated by covarying for total brain volume in all analyses. All data was corrected for multiple comparisons using false discovery rate (FDR) correction.^34^

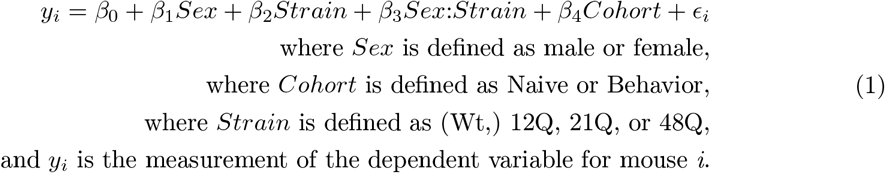

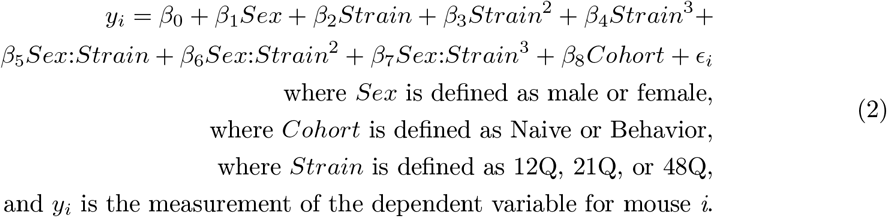

#### 2.2.1 Spatial gene expression analysis

To identify potential mechanisms that might cause these volume changes, we linked spatial gene expression data to the MRI-determined phenotypes. Under the assumption that if a gene is associated with volume change, then subsequent volume changes are spatially restricted to regions in which that gene is expressed, we sought to identify which genes share spatial expression patterns that matched the volume changes associated with *Ar* Q tract lengths.

We used two related gene expression datasets from the Allen Institute to probe this link during neurodevelopment and adulthood: the Allen Developing Mouse Brain Atlas that profiled the expression of 2107 genes related to neurodevelopment over 7 developmental timepoints (E11.5, E13.5, E15.5, E18.5, P4, P14, and P28) and the Allen Mouse Brain atlas that profiled 4345 expression images in adult mice (P56)(limited number of genes due to the use of the coronal dataset only).^35, 36^

Spatial correlation between structural volume and gene expression at each voxel were computed for each gene. Those genes were then ranked by correlation to the *Ar* quadratic phenotype (Equation 2) to assess which genes’ expression is most similar and most dissimilar to the observed volumetric changes. Furthermore, modules (an annotated group of genes (see the Gene Ontology (GO) consortium^37, 38^) that are most and least correlated were extracted. We subsequently used sparse regression to predict spatial volume changes from spatial gene expression. Beyond phenotype prediction, an additional advantage of this method over independent correlations for each gene is the sparse selection of groups of genes.

Detailed methods are described in the Supplementary Methods; these data processing and spatial enrichment methods have been previously used elsewhere, both in exploratory methods,^39, 40^ and to support targeted hypotheses.^41^

### 2.3 Behavioral Phenotype

#### 2.3.1 Statistics

Behavioral data was analyzed using the R statistical language (R Core Team, 2017).^33^ Distributions of the measures were assessed for normalcy and corrected when necessary. The data was assessed using a linear model to determine the strain effect, compared to Wt (see Equation 3). As with the imaging data, the Wt mice were subsequently removed from the analysis and two models of Equation 3 were compared to assess the effect of Q tract length on behavior: one that assumed that repeat length is a linear term (referred to as “linear”), and one that treated the repeat lengths as ordered terms (referred to as “quadratic”). All data was corrected for multiple comparisons using false discovery rate (FDR) correction,^34^ with all p-values pooled together.

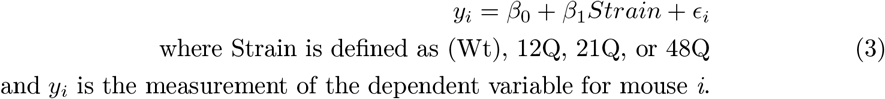

#### 2.3.2 Behavioral Tests

A description of each of the behavioral tests follows, in the order they were performed:^42, 43^

##### Sociability

The three-chambered sociability apparatus consists of a rectangular box with three chambers, with a wire cup placed in either side chamber. The mouse is habituated for 10 minutes during which it can access all three chambers. After this time, a novel stranger mouse is placed under a wire-cup in one of the side chambers. The mouse is observed for an additional 10 minutes during which measures such as time (s) spent in each of the three chambers, specifically near/sniffing the wire cups (near the novel mouse) are observed using an overhead video system, manually scored, and then compared. Mice that are not considered social spend either an equivalent amount of time between the two cups or show a preference for the empty cup (away from the novel mouse).

##### Grooming

For the grooming test, the mice are placed into an empty standard cage (Green Line IVC Sealsafe PLUS Mouse, Techniplast) for ten minutes (habituation). The following ten minutes, mice are placed into a different, empty, standard cage and manually scored for grooming behaviors. Primary measures are duration of grooming bouts (s) as well as number of grooming bouts. Mice with a repetitive behavior phenotype spend significantly more time grooming, with more bouts of grooming, than wildtype controls.

##### Open Field

The open field test consists of 10 minutes of testing, where the mouse is placed in an empty box (area=44cm^2^) and allowed to roam free for the duration of the test. The mouse’s motion is recorded by an automated system (Activity Monitor 7, Med Associates Inc, Fairfax VT) and primary measures investigated are time (s) spent in the center (anxiety) and total distance (cm) moved (hyperactivity).

##### Marble Burying

For the marble burying test, the mice are placed into a standard cage (Green Line IVC Sealsafe PLUS Mouse, Techniplast) with 5cm of bedding. 10 marbles are placed in two columns of five. A photo was taken before testing began, then mice were placed in. After 20 minutes, the number of marbles more than 2/3rds buried was counted, and a photo was taken. A custom-made program in MatLab (The Mathworks Inc, 2013a)^44^ was used to align and compare before and after photos to determine the average distance that the marbles were moved (mm). The test for marble burying is often considered a test of anxiety, with more marbles buried indicative of increased anxiety.

## 3 Results

### 3.1 Altering Q tract length leads to neuroanatomical differences in a non-linear dosage-dependent fashion

The initial imaging analysis (using Equation 1) showed a tendency towards a pattern of decreased volumes in the 21Q brain when compared to wildtype and an increased volume in the 48Q compared to wildtype, with very little difference between 12Q and wildtypes. Specifically, the absolute volume of 66 of the 182 regions were decreased in the 21Q mice, while 43 of the regions were increased in the 48Q mice (see Figure 1A and Figure 2). The overall brain volume of the three mouse models followed the same pattern (see Figure 2A). Though we saw a large proportion of significant sex differences (the absolute volume of 21 brain regions was significantly decreased in females compared to males)(see Figure 1A), no significant strain-sex interactions were observed in any strain. A significant difference was found in the volume of a large proportion of the brain regions between cohorts, with a trend towards increased volumes in the cohort that underwent behavioral testing when compared to the naive cohort (results not shown).

**Figure 1:**
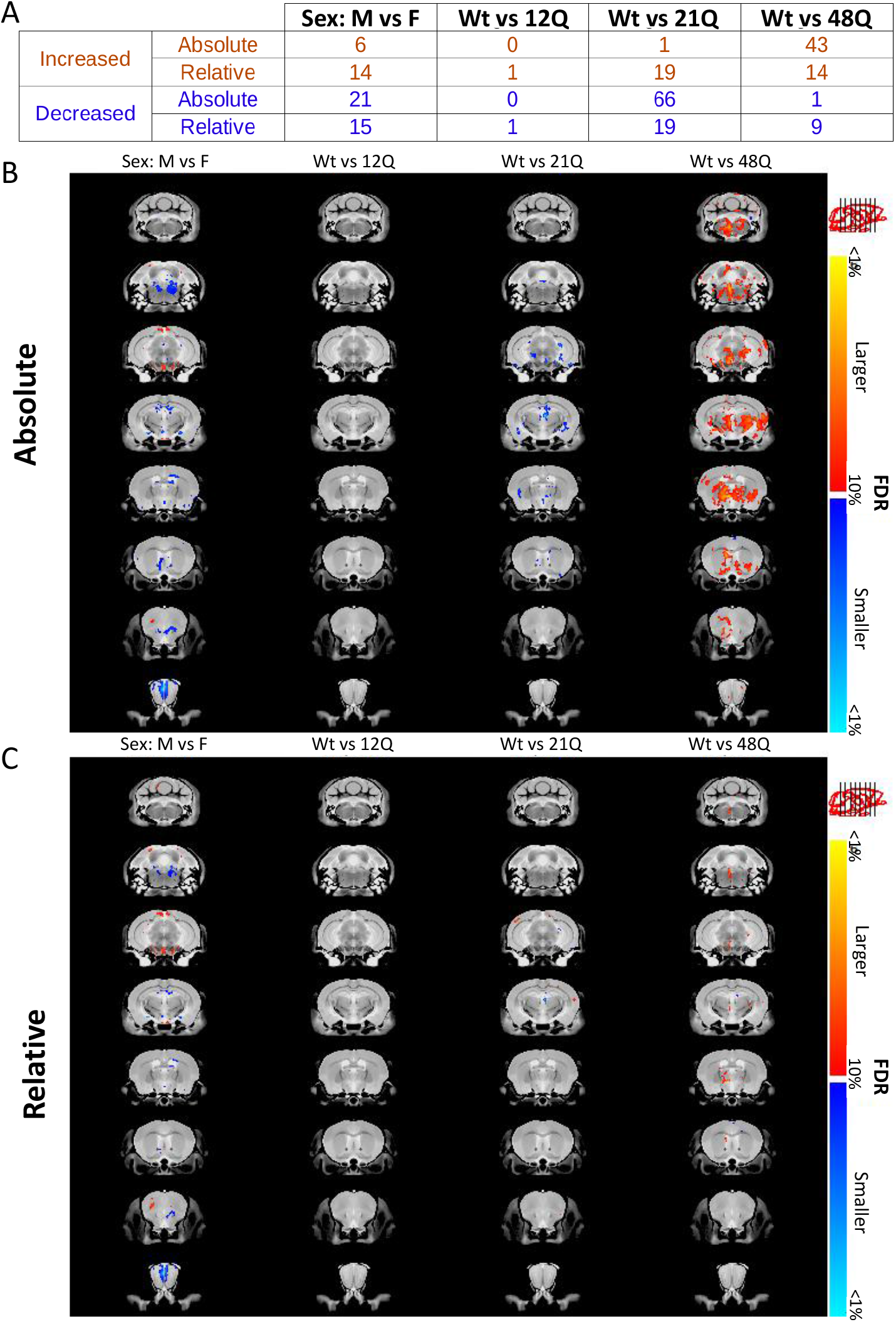
Figure depicting the effect of Q tract length on neuroanatomy (using Equation 1). Table **A** shows the number of regions (from the volume-wise analysis) that were significantly different between groups. Many regions were found to be decreased in the 21Q mouse line, and increased in the 48Q line, while no significant differences were found for the 12Q line. Additionally, no sex-strain interactions were found. Figures **B** and **C** illustrate the voxel-wise changes over the whole brain, with areas with significant increased volume depicted in orange, and decreased volumes depicted in blue.

**Figure 2:**
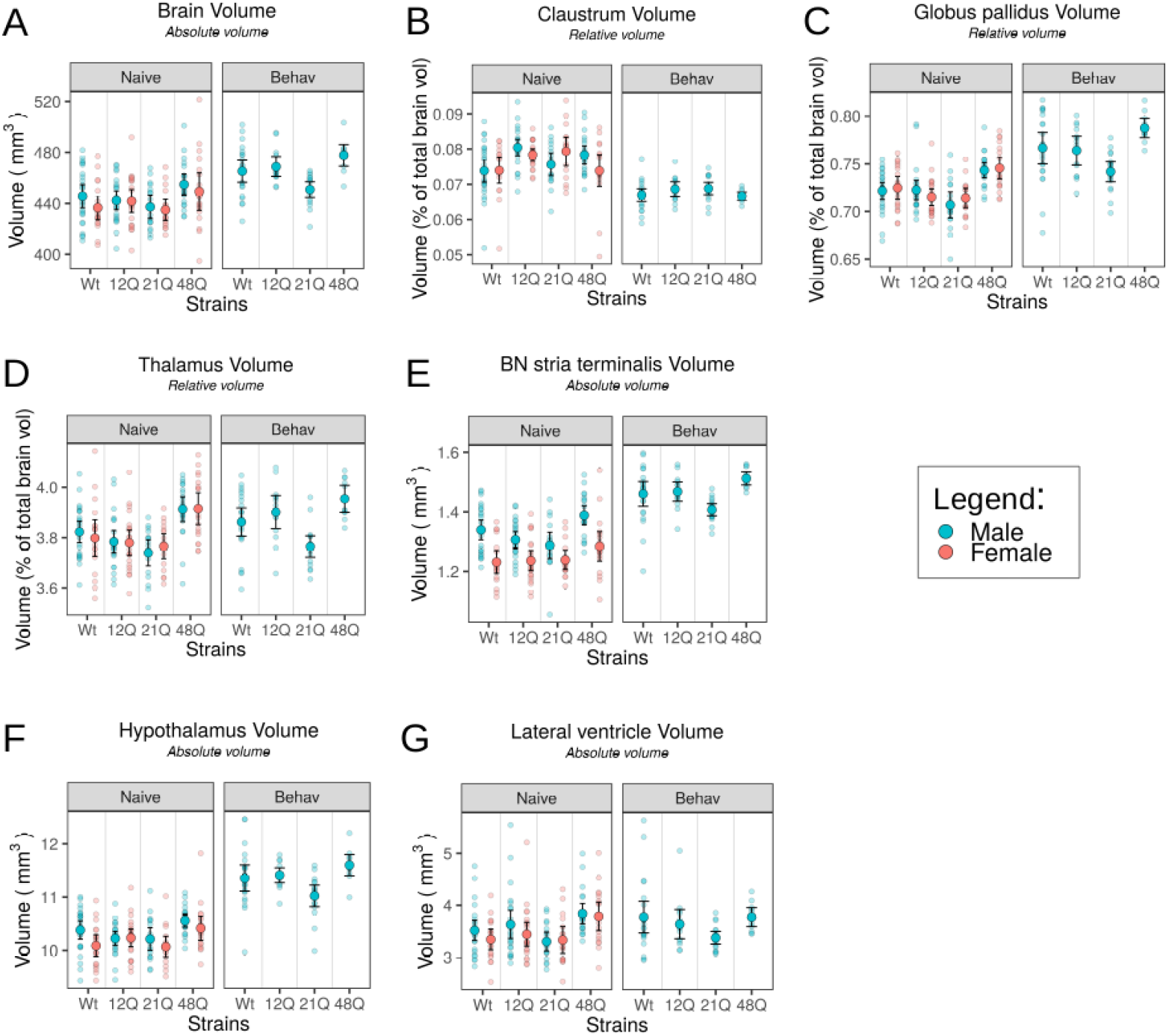
Figures depicting the effect of Q tract length on the volume of the overall brain volume (**A**) and select brain regions (**B**-**G**). Figures **B-D** show the relative volume of the claustrum, globus pallidus, and thamalus as a proportion of total brain volume, while figures **A**, **E-G** show the absolute volume in *mm*^3^. Each point represents one mouse, with blue indicating male and pink indicating female. The error bars represent mean and 95% confidence intervals. The figure for each brain region is faceted to show the results from the naive (“Naive”) cohort as well as the behavior (“Behav”) cohort.

In the subsequent analysis, in which wildtype mice were removed to assess the relationship between the androgen mutant models (see Equation 1 and 2), a non-linear dosage-dependent quadratic relationship was observed, with the 21Q mice showing decreased volumes in more regions compared to the 12Q and 48Q mutants. As before, a large number of sex differences emerged, with no significant strain-sex interactions. Interestingly, the “quadratic” relationship was more pronounced in the absolute volumes of brain regions, while the “linear” relationship was more evident when assessing relative volumes (see Figure 3).

**Figure 3:**
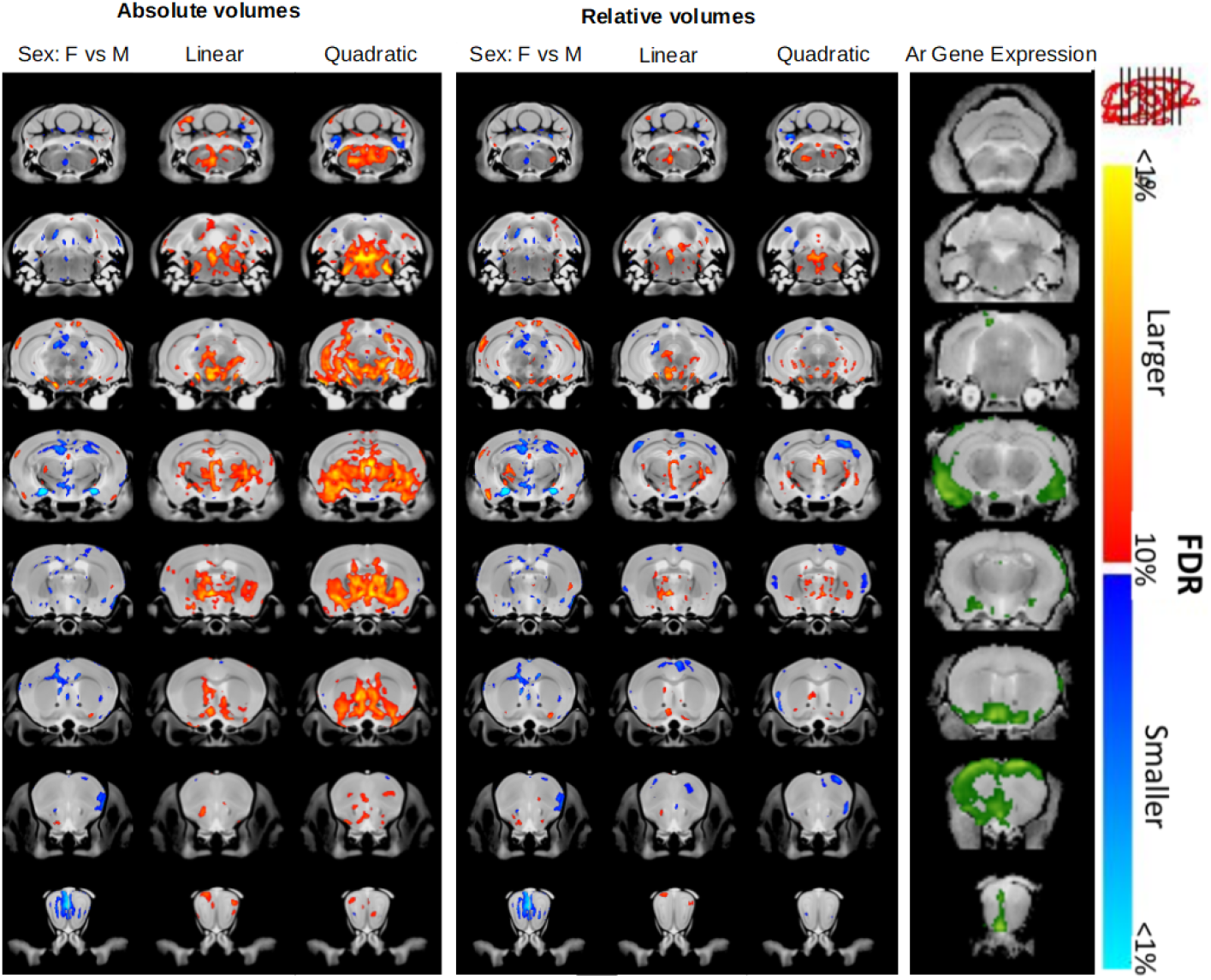
Results from the deformation based morphometry analysis (voxel-wise). Figures depict results from the second analysis, that removed Wt mice to assess the effect of Q tract length on neuroanatomy (see Equations 1 and 2). Significant increases in volume are depicted with hot colours (red-yellow) while decreases are shown with cool colours (blue-torquoise). Columns show sex differences (left), regions where a significant linear pattern was observed (middle)(Equation 1), and regions where a significant quadratic pattern was observed (right)(Equation 2), with absolute and relative volumes differentiated. On the right, *Ar* gene expression information in the adult mouse brain is overlaid on a neuroanatomical map to depict where *Ar* gene expression energy is greatest (green). The most significant differences were observed in the absolute volumes of the quadratic patterns, but did not overlap significantly with the *Ar* gene expression data (fold-change= 0.959). (In contrast, the neurodevelopmental gene expression analysis showed significant overlap with the observed neuroanatomical changes, see Figure 6). Expected sex differences were observed.

### 3.2 Adult neuroanatomical changes due to Q tract length only associated with neurodevelopment

To identify which genes share spatial expression patterns that matched the neuroanatomical volume changes associated with AR Q tract lengths, two related gene expression datasets were used.

To assess the affects of neurodevelopment, LASSO was used to fit a model predicting the spatial AR Q tract length phenotype from spatial expression data of 2107 genes pooled over 7 timepoints. Automated selection of sparsity based on 10-fold cross-validation yielded a subset of 805 images over all 7 timepoints at which the cross-validation error plateaud (see Supplementary data: Figure 1). The selected expression images with the strongest coefficients are related to sex development, and are predominantly images collected at the P28 (adolescence) timepoint. The top 5 genes for both positive and negative coefficients, pooled across all timepoints, are listed in Figure 4. The AR Q tract phenotype and LASSO prediction are shown in Figure 6, along with the spatial expression of genes of interest (see Discussion Section 4.3).

**Figure 4:**
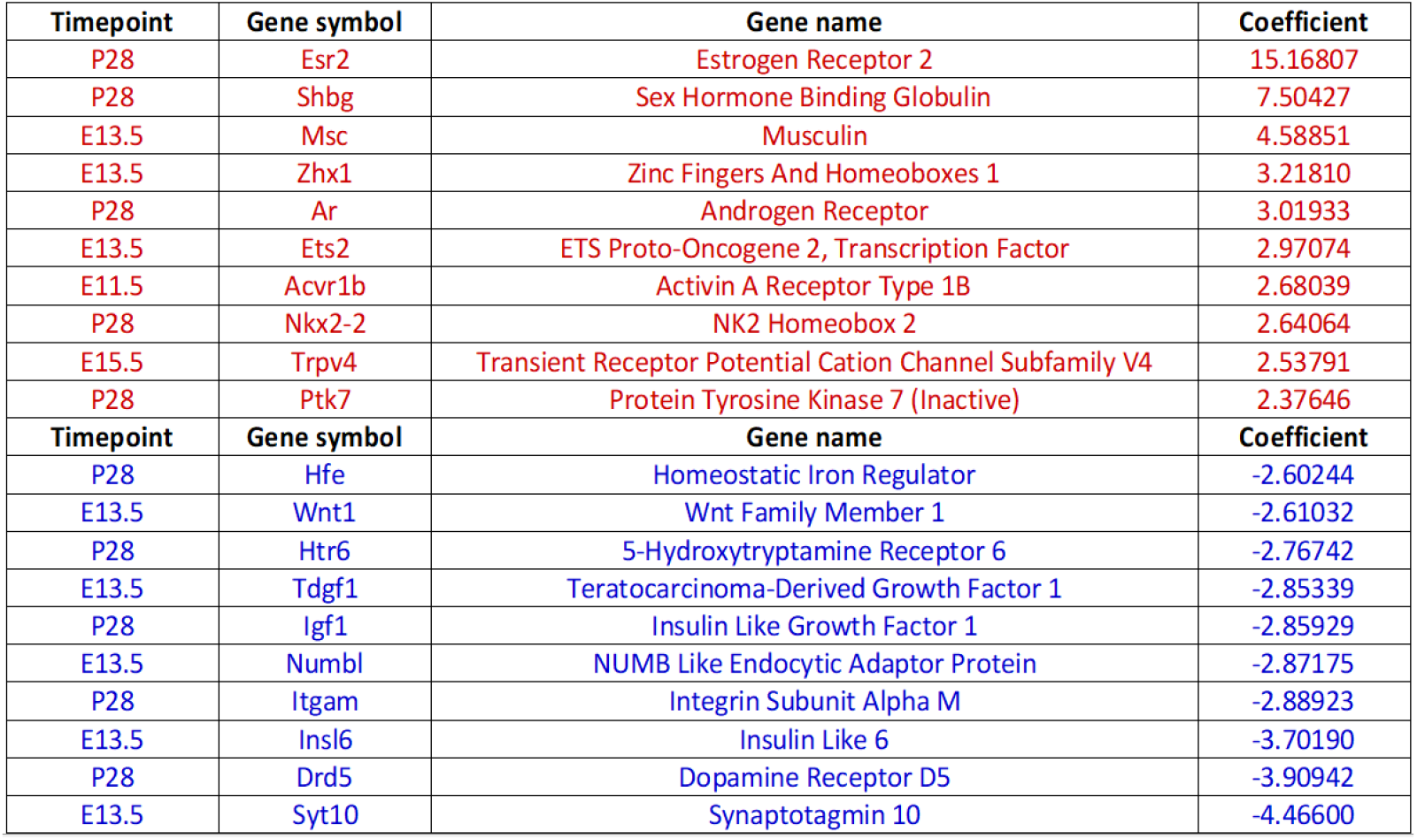
A table of the LASSO coefficients of the five most positively and negatively contributing genes of the AR phenotype, pooled across all time-points. Red indicates positive coefficients and blue indicates negative. The P28 timepoint is associated with the largest coefficients, with Esr2 expression at P28 (which codes for estrogen receptor 2), being by far the most important in predicting the adult AR phenotype. The spatial expression of the *Ar* gene itself at P28 is also an important predictor of the neuroanatomical phenotype

**Figure 5:**
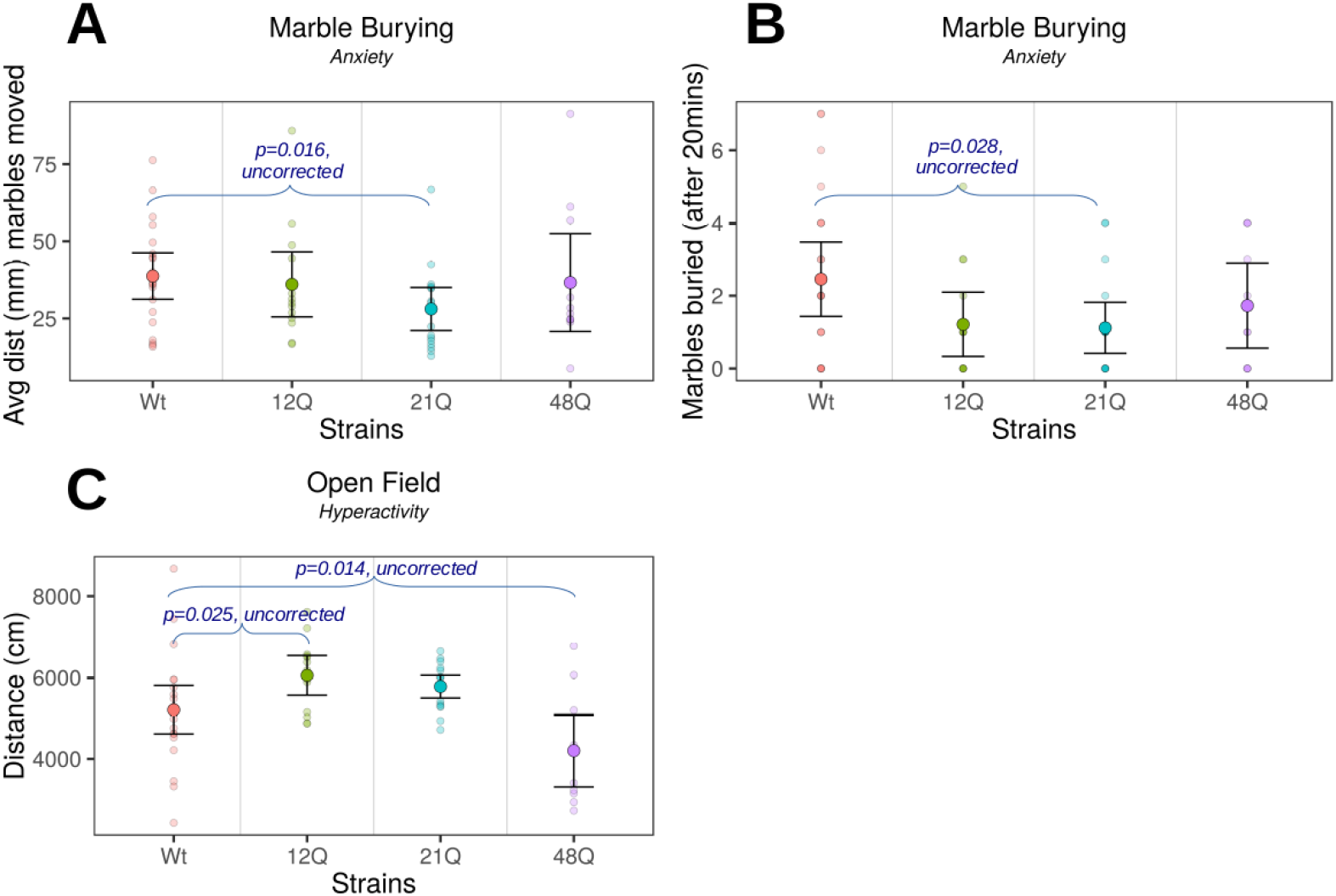
Figures depicting Q tract length effect on the behavior of each strain. Each point represents one mouse. Bars show mean confidence interval (95%). Bars are used to indicate significance levels (not corrected for multiple comparisons), where applicable. Figure **A** shows the average distance (mm) the marbles were moved in the marble burying test, while **B** shows the number of marbles at least 2/3rds buried after 20 minutes. Figure **C** shows the distance (cm) travelled in the open field test.

**Figure 6:**
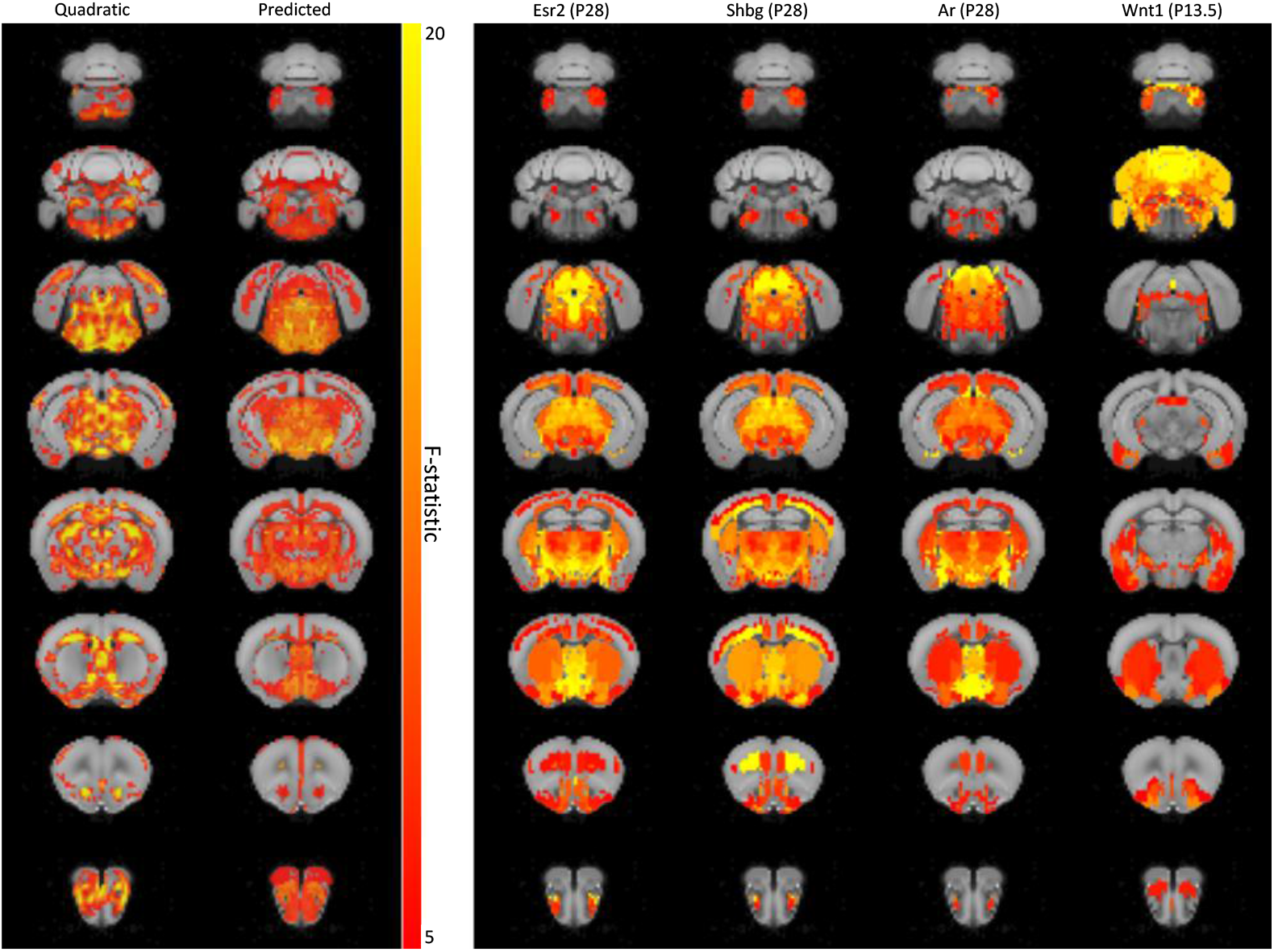
A slice panel showing the LASSO prediction as compared to the AR phenotype. Rows corresponds to various equally spaced coronal slices through the CCFv3 template brain. Shown in each column from left to right are: i)the Ar phenotype from the quadratic analysis (see Equations 2), ii) the LASSO prediction of this phenotype based on spatial gene expression data, and the expression of the following genes: iii) *Esr2* (at P28), iv) *Shbg* (at P28), v) *Ar* (at P28), and vi) *Wnt1* (at E13.5). The colours for the first two columns denote the F-statistic, with more bright regions showing increasingly significant changes in volume. The colours for the gene expression data reflect normalized gene expression (yellow=high, red=low; relative to other regions in the same expression dataset).

Interestingly, the neuroanatomical changes did not correlate with preferential expression of the *Ar* gene of the adult mouse brain. Meaning, there was no greater *Ar* gene expression in the adult mouse brain in regions with neuroanatomical changes that followed the quadratic dosage-dependent pattern (Equation 2)(see Figure 3). None of the genes affiliated with *Ar* had increased gene expression in regions with neuroanatomical differences^45^(Equation 2). Additionally, no module was significantly associated with the observed neuroanatomical changes after correction for multiple comparisons. These results indicate that neurodevelopmental processes, and not gene expression in adulthood, contribute to the neuroanatomical changes observed in adulthood.

### 3.3 No significant effect of Q tract length on behavior in three mouse models

No significant effect of Q tract length was found on any of the behavioral measures, when correcting for multiple comparisons. Without correction, a step-wise effect of androgen sensitivity is seen in the distance travelled (cm) measure of the open field test. The 12Q mice showed significant hyperactivity (p=0.025, uncorrected) and the 48Q line showed significant muted locomotor activity (p=0.014, uncorrected) (see Figure 5). In contrast, both measures of the marble burying test showed decreased anxiety levels (below wildtype) in the 21Q mouse line (p=0.028, p=0.016, both uncorrected). No other measure showed a significant effect of Q tract length, with or without correction for multiple comparisons.

## 4 Discussion

### 4.1 AR activity differentially mediates neuroanatomy and behavior

We expected AR activity to play a linear, dosage-dependent role in both brain and behavior, as well as to mediate sex differences. A general trend was found in the neuroanatomy of the three varying Q tract length mouse models with 21Q mice showing decreased regional volume, 48Q mice showing increased, and no changes in the 12Q. This non-linear but dosage-dependent pattern was unexpected given that the 12Q mice have the greatest amount of AR activity and the 48Q mice have the least.

Moreover, no significant effect of Q tract length was found in the behavior of the three mouse models. When left uncorrected for multiple comparisons, an expected dosage-dependent pattern was observed for the hyperactivity measure, with 12Q mice showing increased hyperactivity and the 48Q mice showing muted locomotor activity. In contrast, the uncorrected analysis of the anxiety measure showed a similar non-linear pattern to the imaging, with 21Q mice showing decreased anxiety when compared to controls. These results indicate that anxiety and hyperactivity are differently and weakly mediated by AR activity, and social behavior and repetitive behavior are not directly mediated by AR activity.

It is interesting to note that in some cases the Q tract length does have a linear relationship with a given trait, while in other cases there may be an optimal length or no effect. These results are not entirely surprising when considering the earlier work investigating these three mouse models. Albertelli et al (2007)^5^ noted that a “simple linear relationship between Q tract length and AR activity may not exist, and different alleles may be optimal in different situations for different reasons.” AR itself has both stimulatory and inhibitory roles, and its primary role is as a tumor suppressor. Moreover, it has been noted that the differences between the lengths is most apparent when context is stressed, such as in limiting the levels of androgen, following castration, or in tumor development.^46^

### 4.2 AR activity alone does not mediate sex differences

Expected sex differences were found in the neuroanatomy of the three strains, but no significant strain-sex interaction effects were found, indicating that sex differences are not mediated by AR activity. Assessing the behavior of female mice with varying Q tract lengths would potentially lead to a better understanding of the mechanisms at play. Henningson et al (2009)^18^ found that only in females was there a significant association between Q tract length and autism, suggesting that perhaps there is a sex difference in susceptibility. Although we found no sex-strain specific interactions in the neuroanatomy, a sex-related effect could be observed in behavioral data if female mutant mice were investigated.

It’s interesting to note the apparent spatial dissimilarity of the sex differences contrast map (Figure 3:“Sex: F vs M”) as compared to the linear and quadratic AR effect maps (Figure 3:“Linear”, “Quadratic”). This is particularly intriguing relative to the debate regarding the role of AR, in contrast to the estrogen receptor, for the masculinization of regional brain anatomy in mice.^47–50^ These results indicate that AR does not contribute significantly to brain morphology, but quantification and further assessment is necessary.

### 4.3 Neurodevelopment processes associated with spatial patterns of volume changes in adulthood

We used two related gene expression datasets from the Allen Institute to probe the link between neuroanatomy and gene expression during neurodevelopment and adulthood. Although there is no overlap of gene expression and volume changes in *Ar* or any of the affiliated genes in the adult mouse brain analysis, the developmental analysis shows timing as an important contributor. Among the top predictors of the AR phenotype are the expression of sex-hormone related genes at P28. Ranked by LASSO coefficient, the *Ar* gene itself is fifth in a list of 14566 predictors, consistent with the idea that affected neuroanatomical regions are those in which the mutated gene is expressed. At P28, *Ar* shares a similar spatial expression pattern to other sex-hormore related genes, including Esr2 (which codes for the estrogen receptor beta) and Shbg (sex hormone binding globulin). Esr2 is by far the most important predictor of this AR Q tract length phenotype. Both the androgen receptor and estrogen receptor beta are downstream targets in the testosterone metabolic pathway.^50^

Of the 7 timepoints that were studied, it is interesting that the strongest predictors are expression maps at P28, the closest timepoint in the expression dataset to puberty (see Figure 4). At P28, these sex-hormone related genes have positive LASSO coefficients, which signifies that regions in which these genes are expressed have an increased volume when Q tract length deviates from the median. The exact mechanism through which hormone-related expression can change brain volumes is unclear, but timing of effect does seem important.

Our spatial gene expression analysis suggests that gene expression at E13.5 might be another mode through which neuroanatomy is affected. Many of the most important predictors at this earlier timepoint are Wnt/beta-catenin signaling related genes, which are implicated in oncogenesis and cell proliferation (see Figure 4. Importantly, beta-catenin is able to directly bind to the androgen receptor in competition with TCF/LEF transcription factors;^51, 52^ the variation in AR activity and resulting brain overgrowth found might be due to increased cell proliferation through this pathway.

Here, we find that Wnt signaling related genes have negative LASSO coefficients, which in this case, suggest that regions in which this pathway is dominant at E13.5 do not increase in volume, in contrast to all other regions (Figure 4). Thus, a potential mechanism that would explain these findings is a global brain overgrowth that is compensated for in the cortex where little volume change is seen. At E13.5, betacatenin and Tcf4 are expressed in most regions of the brain, while *Ar* is expressed in regions that do not show a volume change. Thus, this compensation might occur via Ar/TCF competition for beta-catenin, where increased AR activity results in decreased interactions of beta catenin with TCF/LEF transcription factors, and therefore decreased cell proliferation.

By linking spatial gene expression to neuroanatomical changes, we hypothesize that this AR Q tract length phenotype arises due to two temporally distinct mechanisms mediated through Ar: Wnt signaling during embryonic development and sex-hormone binding during puberty. It should be noted that the methods used here are associative however, and that these potential mechanisms posited post hoc are hypothetical. Experimental work should be done to test these hypotheses. As spatial expression analyses involve correlations over space, they do not take into account actual transcript and protein levels in these mouse models, a fact that must be recognized when interpreting the roles of the recovered genes within signaling pathways. A further limitation is that the developmental gene expression data used—while a fantastic and most comprehensive resource of this type— is limited in coverage across both time (7 timepoints) and the genome (10%).

The results of this study indicate that neurodevelopmental disorder-related behavior and neuroanatomy, as well as sex differences, are likely mediated by more complex mechanisms than AR activity alone. Though this is not in direct contrast to the “extreme male theory” hypothesized in autism,^53^ it does indicate a dependence on more convoluted mechanisms. Given the results of the gene expression data, it is probable that neurodevelopmental processes play a significant role and models that look at other mechanisms are needed to further elucidate what sex-specific mechanisms may be contributing to the disorder and the robust sex-difference observed.

Moreover, although many studies suggested that increased testosterone (and therefore high androgen, as expected in the 12Q mouse model) is associated with autism, some studies suggest the opposite may be true, or that the relationship is not as straightforward as expected. For example, injecting estrogen into the brain of mice lacking reelin improves autism-like features.^54^ Another study found that men with autism have similar testostrone levels as controls but had other less masculine features, while women with autism had increased testostrone and more masculine characteristics.^55^ Also, Henningsson et al (2009)^18^ found that rare mutations in AR are not common in autism, reestablishing the concept that AR activity would likely not be solely responsible for the sex-specific mechanisms underlying autism. To our knowledge, there are no studies assessing the role of AR activity throughout development and the link with autism.

## 5 Conclusion

We investigated the effect of *Ar* Q tract length on the behavior and neuroanatomy of three different mouse lines, in an attempt to elucidate the relationship of AR activity and the phenotype of these mice. The results indicate that androgen sensitivity does not directly lead to autism-related behaviors and neuroanatomy and that the sex-specific mechanisms behind the disorder are more complex, possibly dependent on neurodevelopmental processes. Further study into sex differences, development, other behaviors, and other sex-specific mechanisms are needed to better understand AR sensitivity, neurodevelopmental disorders, and the sex difference in their prevalence.

## 6 Acknowledgements

This work was supported by funding from the Ontario Brain Institute (OBI), the Canadian Institutes of Health Research (CIHR), and Brain Canada. The authors would additionally like to thank Chris Hammill for data analysis assistance, and Christine Laliberte and Dr. Dulcie Vousden for procedural assistance.

