## Supplementary data for "Characterization of Mice Bearing Humanized Androgen Receptor Genes (h/mAr) Varying in Polymorphism Length"

### 1 Supplementary data

#### 1.1 Preferential spatial gene enrichment methods

##### 1.1.1 Gene expression data preprocessing

Spatial gene expression data were downloaded from the Allen Institute's Developing Mouse Brain Atlas via their REST API. These data were profiled for 2107 genes (selected based on their role in neurodevelopment) at 7 developmental timepoints (E11.5, E13.5, E15.5, E18.5, P4, P14, and P28) using ISH. As part of their Informatics Data Processing pipeline, the Allen Institute summarized gene expression within parcellations defined by their Developing Mouse Brain Atlas ontology (ontology ID=12). To allow for cross-timepoint comparisons between the developmental expression data and the adult brain phenotype, we mapped the expression data—in a voxelwise manner—to the Allen Institute's common coordinate framework version 3 (CCFv3) template[1] that describes the adult mouse brain. This was done by obtaining ROI definitions for the Developing Mouse Brain Atlas ontology on this template, and for each gene at each timepoint, setting voxels under each ROI to the appropriate expression value.

We used ANTs (Avants et al., 2008; Avants et al., 2011) to align the MRI template to the CCFv3 template over which the developmental expression data are defined, thereby allowing for voxelwise comparisons between gene expression and neuroanatomy. MRI data was downsampled to 200um to match the resolution of the gene expression data.

##### 1.1.2 Gene expression data analyses

Several approaches were used to investigate the relationship between affected neuroanatomy and spatial gene expression (Allen Brain Institute[2]), in neurodevelopment and adulthood. All analyses were done using the neuroanatomical changes from the quadratic dosage-dependent analysis (see Equation 2).

##### 1.1.3 Gene expression across development

To determine which genes might play a role in shaping this neuroanatomical phenotype, we used the biglasso package (Zeng, 2017) for R (R Core Team, 2018)[3] to fit a LASSO model that predicted the phenotype (a response vector of length 55639 corresponding to voxels under a brain mask) from 14566 predictors (2107 genes at 7 timepoints, with some images excluded due to missing data; predictor matrix size = 55639 x 14566). To determine the optimal sparsity parameter, we used 10-fold cross-validation. This affords the additional advantage (over independent correlations for each genes) of the sparse selection of groups of genes.

##### 1.1.4 Gene expression in adulthood

Preferential spatial expression at a single time-point can be quantified by a fold-change measure (see Fernandes et al 2017[4]): mean gene expression energy in the area of interest (area with neuroanatomical change) divided by the mean gene expression energy in the brain. For every structure, a t-statistic associated with its volume vs Q tract length was computed, along with the structure's mean gene expression energy.

A fold-change value greater than 1 indicates that there is that many times greater expression in regions with neuronatomical differences than the mean expression in the whole brain. This value can be generated for every gene in the adult mouse brain in the Allen Brain Institute.

A mean fold change value was computed for the *Ar* gene, as well as every gene in the Allen Brain Institute. A list of genes associated with *Ar* was created using The Signaling Network Open Resource (SIGNOR)[5] and compared to a list of genes (including *Ar*) that had a fold-change value greater than 1. The mclust R package[6] was used to test whether there was a significant relationship between affected neuroanatomy and spatial gene expression of genes affiliated with *Ar*.

Another method that was used to probe the relationship between neuroanatomy and spatial gene expression in adulthood was to look at the spatial correlation between gene expression and neuroanatomical changes using Spearman's correlation (see Yee 2019 tutorial[7]) and assess associated module/ontology terms (an annotated group of genes (like biological process, synaptic plasticity)(see the Gene Ontology (GO) consortium[8,9]). A list of ranked genes was created, with their associated modules (from GO Biological Processes list[8–10]). Modules were then filtered to only those that contained a minimum of 10 and a maximum of 500 genes. A random phenotype was simulated 5000 times to generate a distribution of AUCs. Pvalues of module association were generated by comparing against the simulation-based AUC distribution and corrected for multiple comparisons, resulting in terms that were most and least associated with the neuroanatomical changes observed.

#### 1.2 Preferential spatial gene enrichment results

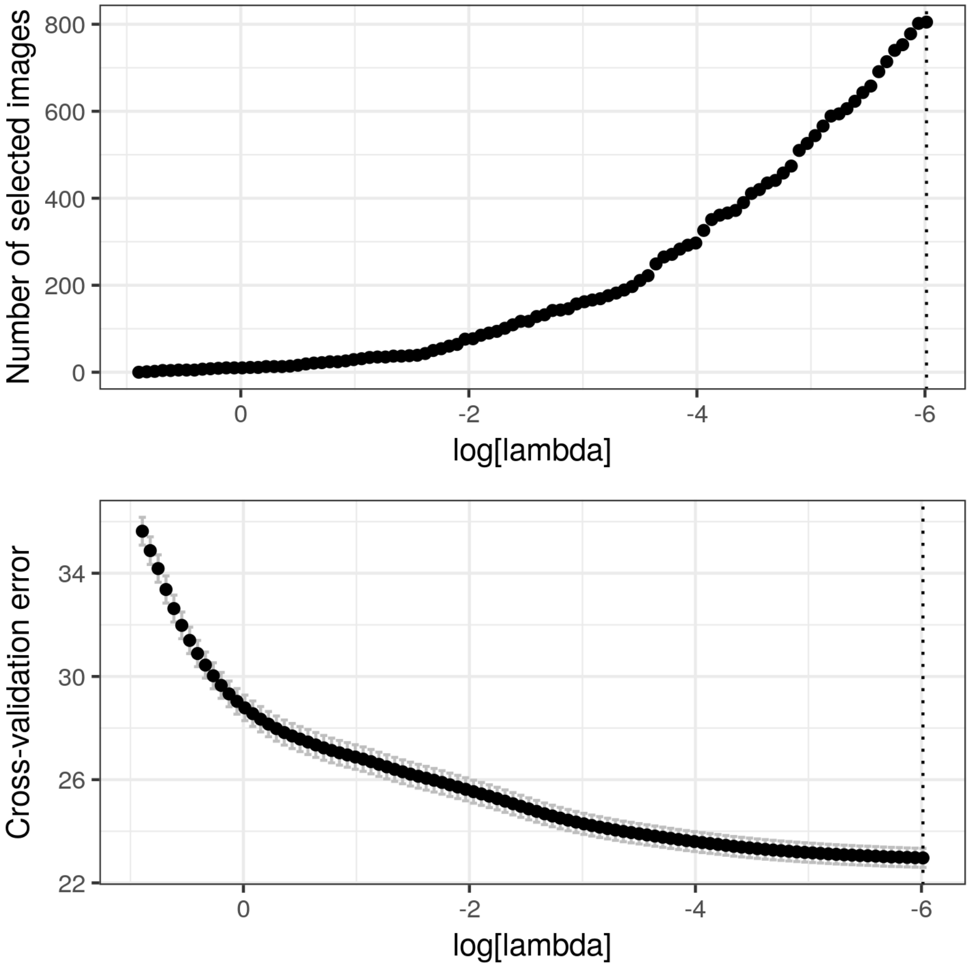


Figure 1: Cross-validation results for the LASSO model. Top panel shows the number of predictors (expression images) as a function of sparsity, bottom panel shows the cross-validation error as a function of sparsity. Error bars shown are standard errors. Automated sparsity selection based on cross-validation suggests a lambda of 0.002445890 ($log[lambda] \sim -6$), at which 805 predictors are selected.

[2] Ed S. Lein, Michael J. Hawrylycz, Nancy Ao, Mikael Ayres, Amy Bensinger, Amy Bernard, Andrew F. Boe, Mark S. Boguski, Kevin S. Brockway, Emi J. Byrnes, Lin Chen, Li Chen, Tsuey-Ming Chen, Mei Chi Chin, Jimmy Chong, Brian E. Crook,

Aneta Czaplinska, Chinh N. Dang, Suvro Datta, Nick R. Dee, Aimee L. Desaki, Tsega Desta, Ellen Diep, Tim A. Dolbeare, Matthew J. Donelan, Hong-Wei Dong, Jennifer G. Dougherty, Ben J. Duncan, Amanda J. Ebbert, Gregor Eichele, Lili K. Estin, Casey Faber, Benjamin A. Facer, Rick Fields, Shanna R. Fischer, Tim P. Fliss, Cliff Frensley, Sabrina N. Gates, Katie J. Glattfelder, Kevin R. Halverson, Matthew R. Hart, John G. Hohmann, Maureen P. Howell, Darren P. Jeung, Rebecca A. Johnson, Patrick T. Karr, Reena Kawal, Jolene M. Kidney, Rachel H. Knapik, Chihchau L. Kuan, James H. Lake, Annabel R. Laramee, Kirk D. Larsen, Christopher Lau, Tracy A. Lemon, Agnes J. Liang, Ying Liu, Lon T. Luong, Jesse Michaels, Judith J. Morgan, Rebecca J. Morgan, Marty T. Mortrud, Nerick F. Mosqueda, Lydia L. Ng, Randy Ng, Geralyn J. Orta, Caroline C. Overly, Tu H. Pak, Sheana E. Parry, Sayan D. Pathak, Owen C. Pearson, Ralph B. Puchalski, Zackery L. Riley, Hannah R. Rockett, Stephen A. Rowland, Joshua J. Royall, Marcos J. Ruiz, Nadia R. Sarno, Katherine Schaffnit, Nadiya V. Shapovalova, Taz Sivisay, Clifford R. Slaughterbeck, Simon C. Smith, Kimberly A. Smith, Bryan I. Smith, Andy J. Sodt, Nick N. Stewart, Kenda-Ruth Stumpf, Susan M. Sunkin, Madhavi Sutram, Angelene Tam, Carey D. Teemer, Christina Thaller, Carol L. Thompson, Lee R. Varnam, Axel Visel, Ray M. Whitlock, Paul E. Wohnoutka, Crissa K. Wolkey, Victoria Y. Wong, Matthew Wood, Murat B. Yaylaoglu, Rob C. Young, Brian L. Youngstrom, Xu Feng Yuan, Bin Zhang, Theresa A. Zwingman, and Allan R. Jones. Genome-wide atlas of gene expression in the adult mouse brain. *Nature*, 445(7124):168–176, 1 2007.

[3] R Core Team. R: A Language and Environment for Statistical Computing, 2017.

[4] Darren J. Fernandes, Jacob Ellegood, Rand Askalan, Randy D. Blakely, Emanuel Dicicco-Bloom, Sean E. Egan, Lucy R. Osborne, Craig M. Powell, Armin Raznahan, Diane M. Robins, Michael W. Salter, Ameet S. Sengar, Jeremy Veenstra-VanderWeele, R.M. Henkelman, and Jason P. Lerch. Spatial gene expression analysis of neuroanatomical differences in mouse models. *NeuroImage*, 163:220–230, 12 2017.

[5] Livia Perfetto, Leonardo Briganti, Alberto Calderone, Andrea Cerquone Perpetuini, Marta Iannuccelli, Francesca Langone, Luana Licata, Milica Marinkovic, Anna Mattioni, Theodora Pavlidou, Daniele Peluso, Lucia Lisa Petrilli, Stefano Pirrò, Daniela Posca, Elena Santonico, Alessandra Silvestri, Filomena Spada, Luisa Castagnoli, and Gianni Cesareni. SIGNOR: a database of causal relationships between biological entities. *Nucleic Acids Research*, 44(D1):D548–D554, 1 2016.

[6] Luca Scrucca, Michael Fop, T Brendan Murphy, and Adrian E Raftery. mclust 5: Clustering, Classification and Density Estimation Using Gaussian Finite Mixture Models. *The R journal*, 8(1):289–317, 8 2016.

[7] Yohan Yee. A practical introduction to spatial gene enrichment analysis in R, 2019.

[8] Michael Ashburner, Catherine A. Ball, Judith A. Blake, David Botstein, Heather Butler, J. Michael Cherry, Allan P. Davis, Kara Dolinski, Selina S. Dwight, Janan T. Eppig, Midori A. Harris, David P. Hill, Laurie Issel-Tarver, Andrew

Kasarskis, Suzanna Lewis, John C. Matese, Joel E. Richardson, Martin Ringwald, Gerald M. Rubin, and Gavin Sherlock. Gene Ontology: tool for the unification of biology. *Nature Genetics*, 25(1):25–29, 5 2000.

[9] The Gene Ontology Consortium. The Gene Ontology Resource: 20 years and still GOing strong. *Nucleic Acids Research*, 47(D1):D330–D338, 1 2019.

[10] Daniele Merico, Ruth Isserlin, Oliver Stueker, Andrew Emili, and Gary D. Bader. Enrichment Map: A Network-Based Method for Gene-Set Enrichment Visualization and Interpretation. *PLoS ONE*, 5(11):e13984, 11 2010.
